# The prevalence and distribution in genomes of low-complexity, amyloid-like, reversible, kinked segment (LARKS), a common structural motif in amyloid-like fibrils

**DOI:** 10.1101/2020.12.08.415679

**Authors:** Michael P. Hughes, Luki Goldschmidt, David S. Eisenberg

## Abstract

Membraneless Organelles (MLOs) are vital and dynamic reaction centers in cells that organize metabolism in the absence of a membrane. Multivalent interactions between protein Low-Complexity Domains (LCDs) contribute to MLO organization. Our previous work used computational methods to identify structural motifs termed Low-complexity Amyloid-like Reversible Kinked Segments (LARKS) that can phase-transition to form hydrogels and are common in human proteins that participate in MLOs. Here we searched for LARKS in proteomes of six model organisms: *Homo sapiens, Drosophila melanogaster, Plasmodium falciparum, Saccharomyces cerevisiae, Mycobacterium tuberculosis*, and *Escherichia coli*. We find LARKS are abundant in *M. tuberculosis, D. melanogaster*, and *H. sapiens*, but not in *S. cerevisiae* or *P. falciparum*. Abundant LARKS require high glycine content, which enables kinks to form in LARKS as is illustrated in the known LARKS-rich amyloid structures of TDP43, FUS, and hnRNPA2, three proteins that participate in MLOs. These results support the idea of LARKS as an evolved structural motif and we offer the LARKSdb webserver which permits users to search for LARKS in their protein sequences of interest.

## Introduction

A new area of cell biology is the study of membraneless organelles (MLOs) in the organization of cellular structures and metabolism. Many MLOs are RNA and protein assemblies that fulfill specific functions for the cell. Examples of MLOs in human cells include P bodies that degrade mRNA, stress granules (SGs) that store mRNA during stresses, and the nucleolus that processes ribosomal rRNA. MLOs also function in other organisms such as germline P granules in *C. elegans* and stress granules in yeast. The aforementioned organelles are not enveloped by membranes to partition them from the cytoplasm but instead organize through multivalent networks of homotypic and heterotypic reversible interactions between proteins and nucleic acids (Li et al., 2012; Alberti et al., 2019). These reversible networks allow MLOs to be dynamic. They may assemble and disassemble in response to stimuli like SGs do in response to stresses, and dissolve as stresses subside. Proteins in MLOs often contain low-complexity domains (LCDs) that help to drive reversible organization (Kato et al., 2012; Kato & McKnight, 2018).

LCDs are regions of proteins with significant biases for one or a few amino acids. An example is the LCD of FUS where four amino acids glycine, tyrosine, serine, and glutamine account for 80% of the LCD composition. The LCD from FUS has also been termed an intrinsically disordered region (IDR) or a prion like domain (PrLD). The LCD of FUS is an IDR because for most of the time it lacks a defined globular structure. In fact, LCDs are a reasonable proxy for IDRs (Wootton, 1994), but this is not always the case (e.g. collagen proteins which are low in sequence complexity but have a defined poly-proline type II structure). PrLD technically refers to a domain that resembles a yeast prion sequence that is rich in asparagine and glutamine (Cascarina et al., 2017). Hence yeast prions are low-in complexity by definition and PrLDs are typically disordered until forming amyloid fibrils. FUS was identified as a PrLD in humans by a search for proteins with sequence biases similar to yeast prions, alongside a number of other proteins that are involved in SGs including hnRNPA1, hnRNPA2, and TDP43 (Kato et al., 2012. Cell; Molliex et al., 2015. Cell; Lin et al., 2015; Cascarina et al., 2017). However not all LCDs are PrLDs; Arginine-GLycine-rich LCDs are common in RNA binding proteins and are less prone to aggregate than other LCDs (Schwartz et al., 2013).

Remarkably, when the LCDs from hnRNPA1, hnRNPA2, FUS, and TDP43 are purified they undergo liquid-liquid phase-transition and eventually form hydrogels composed of amyloid-like fibrils (Kato et al., 2012; Molliex et al., 2015; Patel et al., 2015; Hennig et al., 2015; Murakami et al., 2015). Phase transitions are also the governing force forming MLOs, and some proteins contribute to MLO organization through their IDRs and LCDs (Protter et al., 2018). The first phase-transition is a liquid-liquid phase separation that leads to a protein rich phase compared to the surrounding bulk solvent. Some proteins may undergo a second phase transition from liquid to solid to form amyloid fibrils (Patel et al., 2015; Murakami et al., 2015; Vogler et al., 2018; Hughes et al., 2018). hnRNPA1, hnRNPA2, FUS, and TDP43 have all been found aggregated in amyloid diseases and MLOs have been proposed as a crucible for driving fibril formation in ALS (Li et al., 2013; King et al., 2012; Shin & Brangwynne et al., 2017).

Amyloid fibrils have long been associated with disease and neurodegeneration, but there are now abundant examples of functional amyloid. Examples include curli fibrils made by *E. coli* to construct biofilms, prions in yeast to alter phenotypes, and PMEL granules in humans to make pigment (Otzen & Riek, 2019; Liebman et al., 2012). In all these examples, the organism takes advantage of the ability of proteins to form fibrils based on mated β-sheets. The fibrils formed from the LCDs of FUS, hnRNPA1, and hnRNPA2 are notable because these amyloid-like fibrils are labile (Kato et al., 2012; Murakami et al., 2015; Hennig et al., 2015) and are easily reversed by elevated temperature, changes of solvent conditions, or dilution. This lability enables amyloid fibrils formed by LCDs to participate in the organization and dynamics of MLOs, but contrast with the detrimental stability of pathogenic amyloid.

In previous work, we identified short adhesive motifs termed Low-complexity Amyloid-like, Reversible Kinked Segments (LARKS) that capture the reversible fibril behavior of these proteins (Hughes et al., 2018). LARKS allow proteins to form a β-sheet rich structures that hydrogen bond along the fibril axis to create amyloid-like fibrils, but the LARKS introduce sharp kinks in the peptide backbone which interrupts the pleated β-sheets (Figure 4A). This structure contrasts with the adhesive elements we find in amyloid fibrils called steric zippers that form extended, pleated β-sheets with interdigitated sidechains giving amyloid fibrils their typical stability (Figure 4B) (Sawaya et al., 2007). The LARKS structure is stable enough to form a fibril, but avoids the irreversibility of a steric zipper, and appears to function in the organization of MLOs.

We computationally searched for LARKS motifs in the human proteome by threading protein sequences onto known LARKS structures and using a Rosetta energy algorithm to predict if each segment can adopt a LARKS structure (Goldschmidt et al., 2010; Hughes et al., 2018). This search through the human proteome found that LARKS are common in LCDs of proteins found in MLOs. Here, we extend this search for LARKS in the organisms *Escherichia coli, Mycobacterium tuberculosis, Saccharomyces cerevisiae, Plasmodium falciparum, Drosophila melanogaster* to compare to the distribution of LARKS in the *H. sapiens* proteome. These model organisms were chosen because they are well studied and cover an array of complexity. *E. coli* and *M. tuberculosis* are both prokaryotes, but *M. tuberculosis* is an intracellular parasite. *P. falciparum* was chosen as a eukaryotic parasite, *S. cerevisiae* as a single celled eukaryote, and *D. melanogaster* as an example multicellular eukaryote. We compare the LARKS predictions for these model organisms to make several findings.

Threading reveals that not all species have LARKS-rich LCD domains, as the LCDs of *S. cerevisiae* and *P. falciparum* are not enriched in LARKS. This correlates with a lack of glycine in their LCD amino acid bias. We go onto provide examples of how LARKS and amino acid composition influences amyloid structure. Overall, this work helps us to understand the roles that LARKS play in biology. We make all our LARKS predictions publicly available with our online database LARKSdb (http://servicesn.mbi.ucla.edu/LARKSdb/).

## Results

### LARKS-rich proteins overlap with LCD-containing proteins in most species

Past work found that in the proteins with the most LARKS overlapped with LCD-containing proteins (Hughes et al, 2018). Here we have reconfirmed this result by sorting the human proteome in two ways: the proteins with the most LARKS per 100 residues and another list of the proteins with the greatest fraction of LCD content. We compared the intersection of the top 5% of proteins from each list and found a large overlap between them of 27% in *H. sapiens* (Supplemental Figure 1) in line with previous observations (Hughes et al., 2018). We term proteins that are LARKS-rich and have an LCD domain LARKS∩LCD proteins. Many of these LARKS∩LCD proteins were found in *M. tuberculosis* (51%) and *D. melanogaster* (32%). A smaller intersection of these two populations was found in *E. coli* (14%), *S. cerevisiae* (17%), and *P. falciparum* (4%) indicating that in these species the proteins with the most LARKS do not overlap with LCD-containing proteins. This preliminary look foreshadows the variability of LARKS content among LCDs in different organisms.

We hypothesized that LARKS are an enriched motif across LCDs of organisms that use them to organize MLOs, as in *H. sapiens*, and searched our threading data to find LARKS-rich proteins. We defined a LARKS-rich protein as any protein having more LARKS per 100 residues than the average value for that species’ proteome. We found that most, but not all, LCD-containing proteomes are rich in LARKS. In *H. sapiens*, we see that 47% of the LCD-containing proteins are LARKS-rich while only 36% of proteins without LCDs were considered LARKS-rich (Figure 1). This indicates enrichment, and in fact P values from bootstrapping confirmed that LCD-containing proteins in *H. sapiens* are significantly more likely to be LARKS-rich than if the proteins did not have an LCD: P value = 1.0*10^-4 (Supplemental Figure 2). Next, we compared LARKS enrichment in LCD-proteins from *H. sapiens* to other proteomes. *E. coli* was chosen as a model bacterial organism with a minimal genome that has very few LCDs. In fact, only 4.4% of proteins have an LCD, but even that small sample is still significantly enriched in LARKS (P value = 3.0*10^-4) (Figure 1; Supplemental Figure 2). To compare to another higher multicellular organism, we chose *D. melanogaster* and find it has similar LCD and LARKS content to *H. sapiens. D. melanogaster* LCD containing proteins are also enriched in LARKS (P value = 3.0*10^-4). Next, we compared the *S. cerevisiae* proteome which is a single celled eukaryote known for its extensive LCDs that form prions. 17.6% of the *S. cerevisiae* genome has LCDs, and to our surprise they are not significantly enriched in LARKS (P value = 0.28). LARKS are a structural motif abundant in LCD-containing proteins of *H. sapiens* and *D. melanogaster*, but not in *S. cerevisiae*.

**Figure 1.**
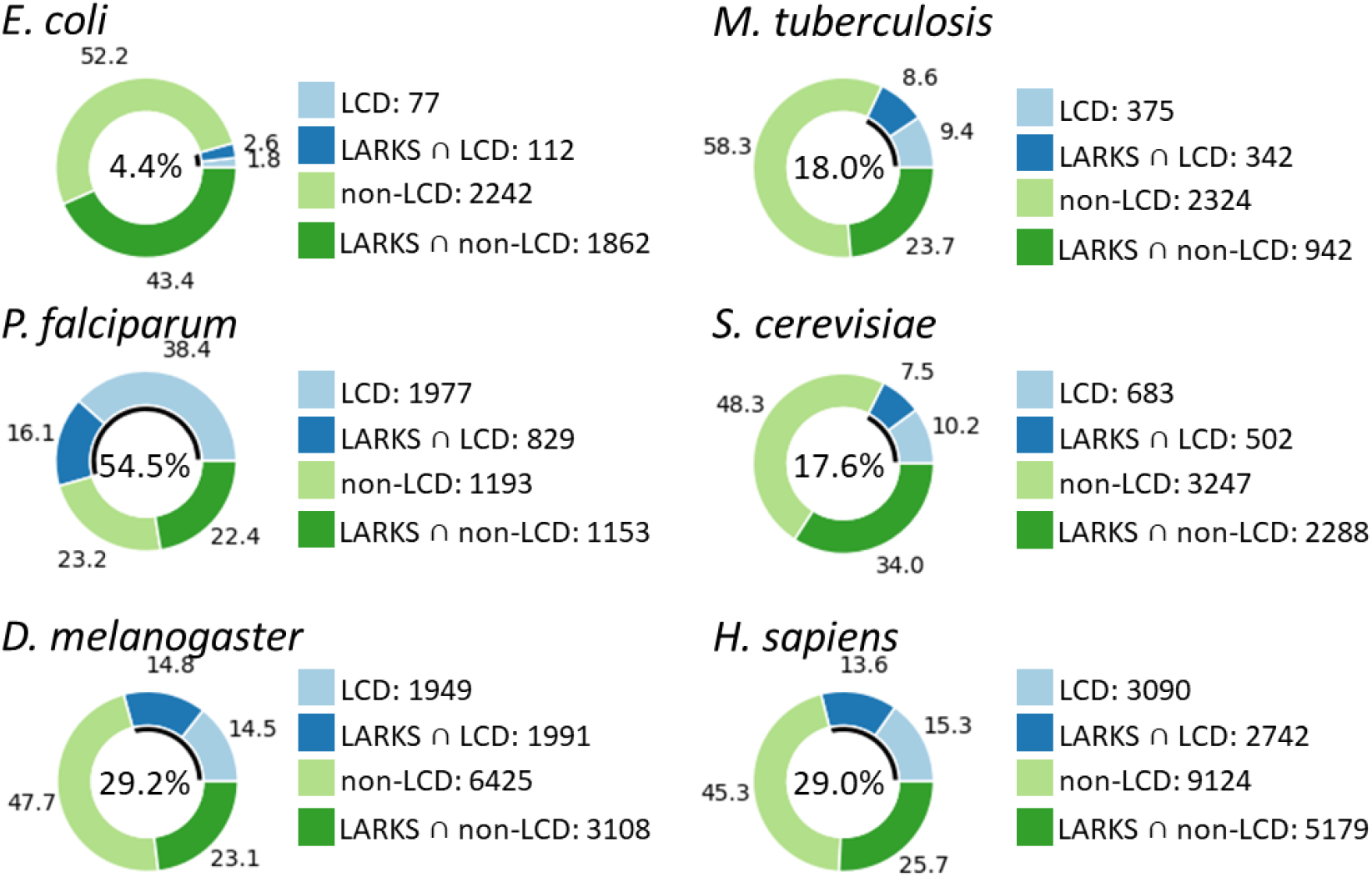
LARKS and LCDs of proteins in six analyzed proteomes. The center of each donut chart gives the percentage of proteins in the proteome that have an LCD and is represented by the black arc in the inside of the donut chart. The outer ring of the donut chart shows the percentage of proteins that are LARKS-rich with an LCD (Dark blue: LARKS ∩ LCD), just LCD-containing (Light blue: LCD), LARKS-rich without having an LCD (Dark green: LARKS ∩ non-LCD), or non-LCD-containing (Light Green: non-LCD). The integers listed next to the keys give precise values for the number of proteins in each category for their respective proteomes.

Following this we compared other single celled organisms with substantial LCD content and chose the two intracellular obligate parasites: the *M. tuberculosis* bacterium responsible for tuberculosis and the *P. falciparum* eukaryote responsible for malaria. *M. tuberculosis* has a proteome where 18% of proteins have an LCD, and LARKS-rich proteins are significantly enriched in the LCD containing proteins (P value = 1.0*10^-4). The opposite was found with *P. falciparum* where 54.5% of proteins contain LCDs and of those LARKS-rich proteins are significantly underrepresented (P value = 1.0*10^-4). Contrary to the LARKS∩LCD-rich proteomes of *E. coli, M. tuberculosis, D. melanogaster*, and *H. sapiens* we consider *S. cerevisiae* and *P. falciparum* to have LARKS∩LCD-poor proteomes. In summary, the analysis shows that proteomes differ markedly in their proportion of LARKS∩LCD proteins.

### LARKS residues overlap with LCDs in LARKS-rich proteomes

The above analysis found the extent to which LARKS motifs are included in whole proteins that contain an LCD. Next, we examine the extent to which the LCDs of proteomes are enriched in LARKS. We also examine the extent to which proteins contain LCDs where LARKS do not reside in the LCD. To do so, we summed the residues of each proteome and counted the number of residues that are in both an LCD and a LARKS (Table 1) and called these LARKS∩LCD residues.

**Table 1.** Residue content of proteomes. The percent of residues in each proteome that were found to be in LARKS, LCDs, or globular regions are shown in columns 4-6. LARKS ∩ LCD Actual gives the percent of residues that are LARKS in LCDs. The expected intersection value is the fraction of LARKS residues multiplied by the fraction of LCD residues. The same methodology is repeated to find LARKS ∩ Glob and LCD ∩ Glob. Comparing the actual intersection of categories with the expected indicates that LARKS are depleted in globular regions in all proteomes and that LARKS residues are more likely to be in LCDs in all organisms studied except *P. falciparum*.

| | Number proteins | Number LCD proteins | % LARKS | % LCD | % Glob | % LARKS ∩ LCD | | % LARKS ∩ Glob | | % LCD ∩ Glob | |
| --- | --- | --- | --- | --- | --- | --- | --- | --- | --- | --- | --- |
|  |  |  |  |  |  | Actual | Expected | Actual | Expected | Actual | Expected |
| E. coli | 4293 | 199 | 2.34 | 0.61 | 85.55 | 0.02 | 0.01 | 1.82 | 2.01 | 0.43 | 0.52 |
| M. tuberculosis | 3983 | 717 | 3.88 | 5.35 | 75.24 | 1.25 | 0.21 | 1.91 | 2.92 | 1.7 | 4.03 |
| P. falciparum | 5152 | 2806 | 0.81 | 11.21 | 77.74 | 0.09 | 0.09 | 0.49 | 0.63 | 4.63 | 8.72 |
| S. cerevisiae | 6720 | 1185 | 1.64 | 2.43 | 77.37 | 0.06 | 0.04 | 1.05 | 1.27 | 0.97 | 1.88 |
| D. melanogaster | 13473 | 3940 | 2.24 | 5.63 | 71.84 | 0.35 | 0.13 | 1.07 | 1.61 | 2.01 | 4.05 |
| H. sapiens | 20135 | 5832 | 2.07 | 3.83 | 70.56 | 0.18 | 0.08 | 1.12 | 1.46 | 1.22 | 2.7 |

We compare this number to the expected probability of this intersection calculated from the fraction of LCD residues multiplied by the fraction of LARKS residues. If the actual counted intersection of LARKS∩LCD residues is greater than expected, we consider the LCDs to be enriched in LARKS.

Overall, these results show that for *E. coli, M. tuberculosis, D. melanogaster*, and *H. sapiens* the actual number of residues that intersect between LARKS and LCDs is over twice that expected by probability (Table 1). This agrees with the previous analysis that LARKS are enriched in LCDs, and that the LARKS reside within the LCD residues in these organisms. *S. cerevisiae* shows a slight enrichment, and *P. falciparum* is about the same as expected. This largely agrees with the data from Figure 1 and goes further by showing that LARKS reside in the actual LCDs of organisms that have LARKS∩LCD proteins.

We searched for LARKS in LCDs because LCDs are typically disordered, and LARKS need to be solvent exposed to allow interactions that affect function. While LCDs are a reasonable proxy for disorder, they are not always perfect (Wootton 1994). Therefore, we also repeated the same analysis but looked for predicted globular and disordered regions as defined by the algorithm Glob (Linding et al., 2003). We call LARKS in predicted globular regions LARKS∩GLOB and for each proteome the actual number of LARKS∩GLOB is lower than expected in predicted globular regions (Table 1). This indicates enrichment of the LARKS motifs in IDRs of proteins. In fact, even in proteomes that lack LARKS∩LCD proteins we see that LARKS∩GLOB residues are rarer than expected. We interpret this to mean LARKS are excluded from globular regions indicating the motif preference to be in IDRs regardless if that IDR is an LCD or not. To gain insight into why LCDs in some proteomes are enriched in LARKS while others are not, we looked at LCD amino acid compositions of proteomes.

### Amino Acid biases are different depending on species

To gain understanding of the variation among proteomes of the extent of LARKS in LCDs, we examined the amino acid composition of LCDs in different proteomes. We found Glycine to be the most common residue in predicted LARKS. This abundance of glycine is consistent with LARKS structures where the allowed phi-psi angles form kinks in the LARKS while maintaining hydrogen bonds along the fibril axis (Hughes et al., 2018). In the proteomes with LARKS-rich LCDs (*E. coli, M. tuberculosis, D. melanogaster*, and *H. sapiens* (Figure 1) we find that Glycine is among the five most common residues (Figure 2). Asparagine is not one of the five most common residues in these proteomes but is among the top five of the *S. cerevisiae* and *P. falciparum*. The presence of glycine in the top 5 residues amino acids of LARKS∩LCD-rich proteomes juxtaposed to its absence in LARKS∩LCD-poor proteomes is striking. To gain functional insight into the function of LARKS in LCDs we studied amyloid structures from LARKS-rich and LARKS-poor proteins (Discussion).

**Figure 2.**
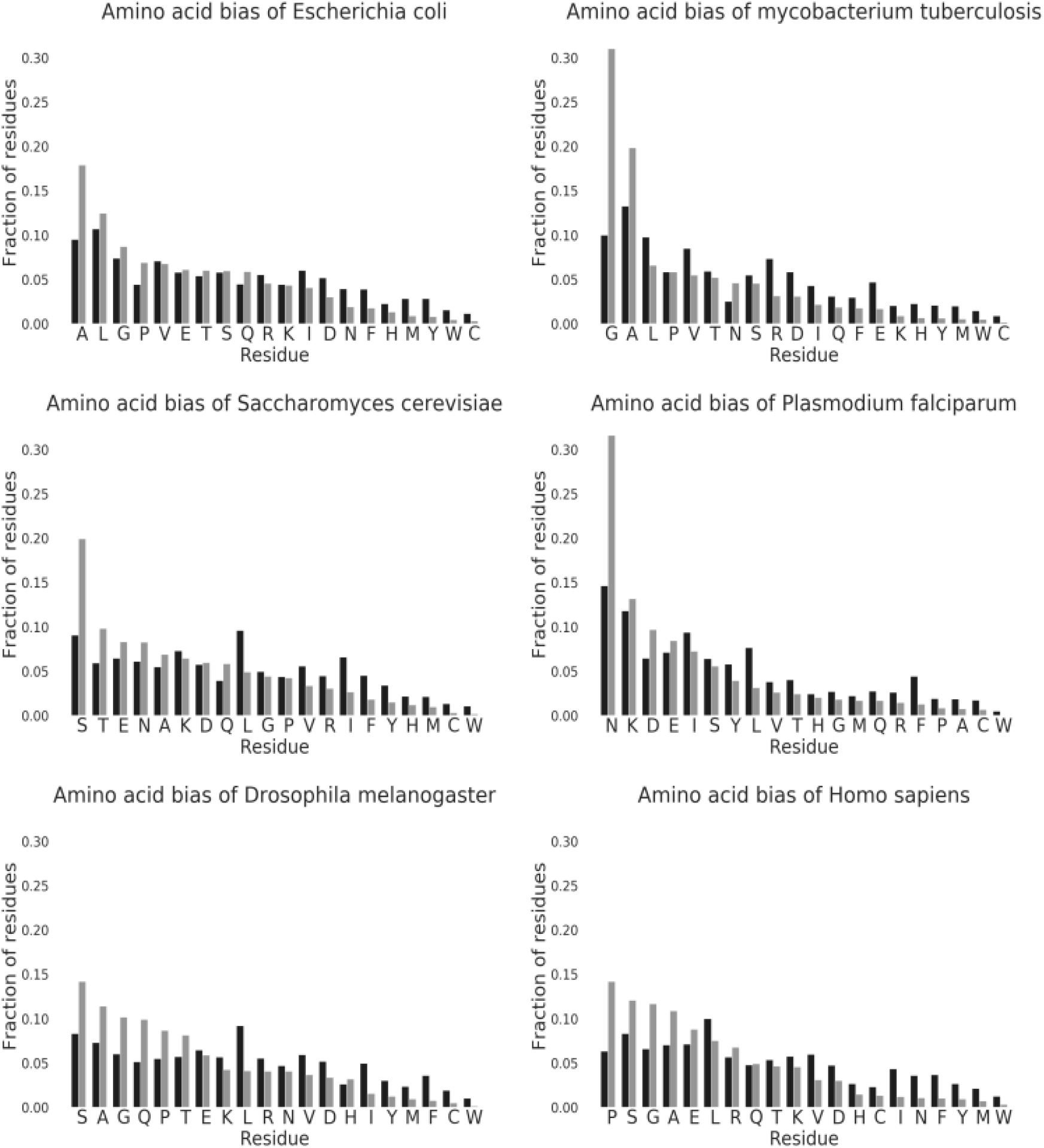
Amino acid bias of proteomes. Black bars represent the fraction of each proteome made up of that amino acid. Gray bars represent the fraction of LCD residues within the proteome made up by each reissue. Amino acids are ordered by the most common amino acids found in that proteome’s LCDs.

## Discussion

LARKS are structural motifs that mediate labile interactions that form reversible amyloids and are associated with phase transitions. Here we use more rigorous methodology to confirm our previous finding that LARKS are motifs enriched in LCDs and IDRs in *H. sapiens*. However, when we applied this same analysis to other organisms, we found a broad spectrum of LARKS content in the LCDs of various species. Across all proteomes analyzed, LARKS are enriched in IDRs (Table 1) indicating that the sharp kink inherent to LARKS is poorly accommodated in globular structure. However, LARKS are not enriched in the LCDs of all proteomes (Figure 1; Supplemental figure 2) and we were curious if the presence of LARKS-rich proteins correlates with biology of the respective species’ proteome as judged by a qualitative analysis of the GO terms over-represented in the LARKS-rich proteins of each genome (Supplemental Table 1).

Of other proteomes we analyzed, *D. melanogaster* is most similar to *H. sapiens* in LARKS abundance and LCD content, and closest to *H. sapiens* in evolutionary distance. GO terms are very similar for LARKS∩LCD proteins between the two organisms. Both proteomes are enriched for proteins involved in RNA binding, transcription factor binding, membaneless organelles (e.g. omega speckles, SGs, nuclear specks, and Cajal bodies), and extracellular matrix/cuticle formation (Figure 5; Supplemental Table 1). In our 2018 analysis of LARKS in the human proteome we identified that keratins are highly enriched in LARKS and posited that phase-separation may be an important aspect of cuticle formation (Hughes et al., 2018) and this was subsequently confirmed in later studies (Quiroz et al., 2020). GO for LARKS∩LCD proteins in *D. melanogaster* included a term for “structural constituent of the chitin-based cuticle” which is the fly barrier equivalent. Chitin is a main component of the *D. melanogaster* exoskeleton, and absent in *human skin*. It will be interesting to see if arthropods use LARKS akin to *H. sapiens* to in cuticle formation.

**Figure 3.**
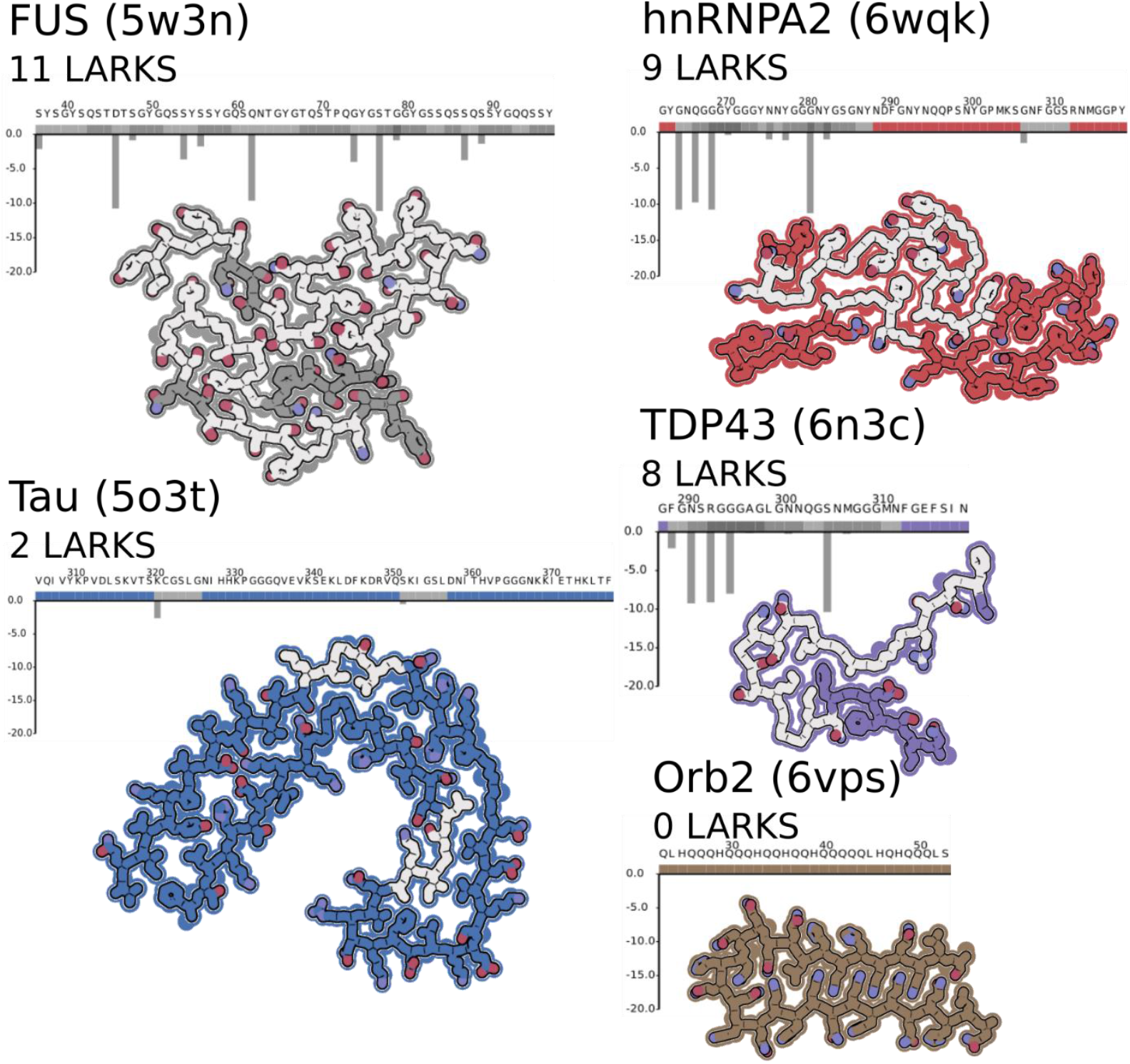
ARKS alter amyloid structure. All images are made from a single chain and a single layer of the fibril, being viewed down the fibril axis. LARKS predictions and sequences are shown above structures. Gray boxes above the sequence indicate predicted LARKS, and the corresponding residues in the protein images have a gray interior. There are three structures of LARKS-rich LCDs: from FUS, hnRNPA2, and TDP43 (PDBIDs: 5w3n, 6wqk, and 6n3c). And two structures from irreversible amyloid Tau and Orb2. Tau is associated with Alzheimer’s pathogenesis and the presented structure (PDBID: 5o3t) was made by seeding purified protein with extracts from the brain of a deceased patient with Alzheimer’s Disease. Orb2 (PDBID: 6vps) from *D. melanogaster* that forms stable amyloid fibrils associated with memory formation in flies and was purified from fly brains. Qualitatively, the structures of proteins associated with dynamic MLOs (FUS, hnRNPA2, and TDP43) have more LARKS and have more kinks in their backbones when compared to the stable amyloids – disease associated (Tau) or functional amyloid (Orb2).

**Figure 4.**
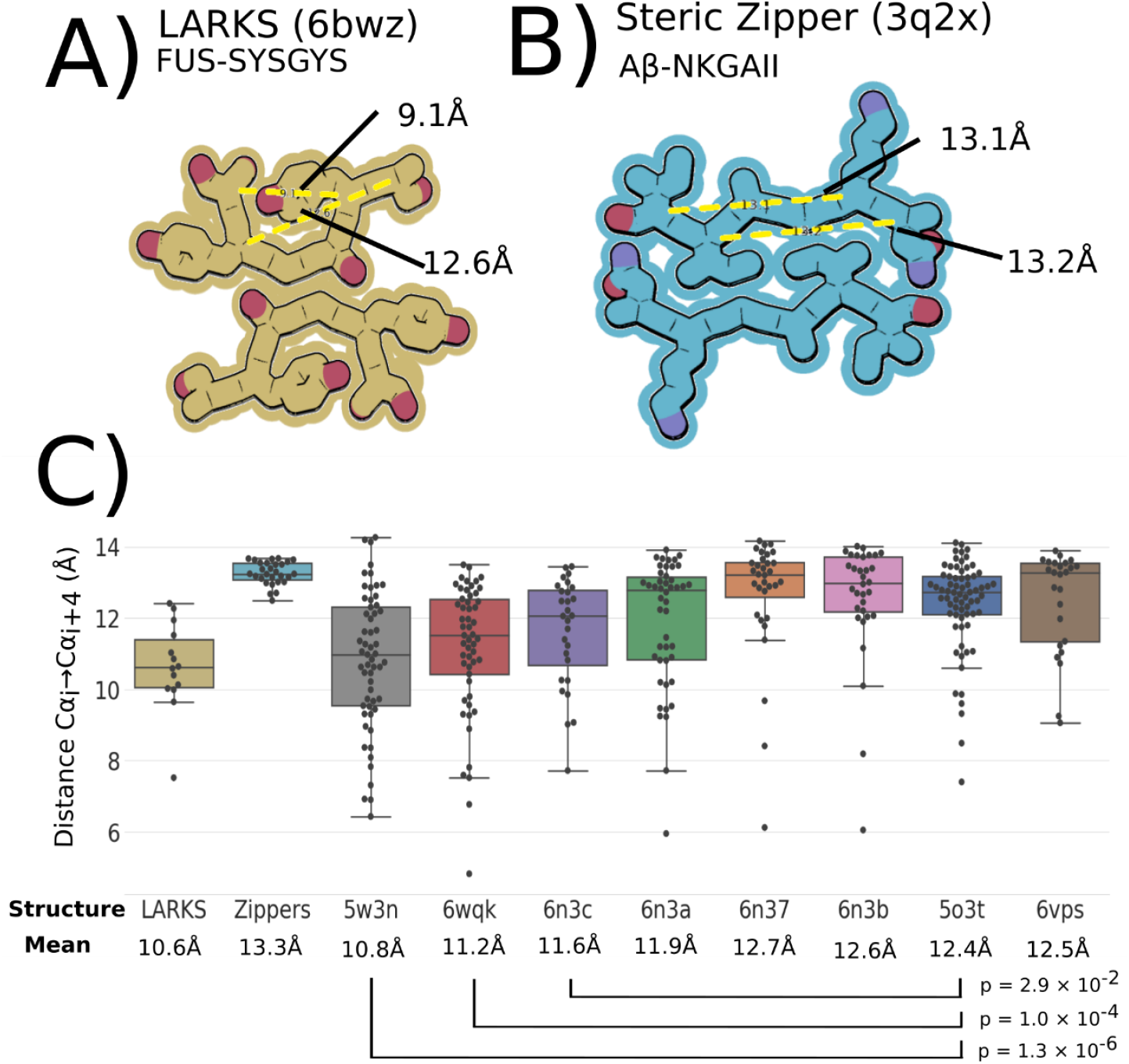
LARKS in protein structure. A) Example of a LARKS structure, FUS segment SYSGYS, shows the kinked backbone and minimal interface between mating sheets. Measuring all possible Cα_i_ →Cα_i+4_ pairs finds distances of 12.6 and 9.1Å, a result of the kinked backbone. B) An example of a steric zipper Aβ-NKGAII to show the extensive mating interface between the extended, pleated β-sheets. Cα_i_ →Cα_i+4_ distances are 13.1 and 13.2Å because the regular spacing from the β-sheet structure. C) Box and whisker plots of Cα_i_ →Cα_i+4_ distances. Each black point is a single measurement from either a sample of LARKS crystal structures, steric zipper structures, and the full-length proteins shown in Figure 3 as well as additional TDP43 structures from Supplemental Figure 3. The LARKS-rich LCD amyloids (FUS-5w3n, hnRNPA2-6wqk, and TDP43-6n3c) have significantly closer Cα_i_ →Cα_i+4_ distances than found in irreversible amyloids (Tau-5o3t and Orb2-6vps) reflecting how LARKS affect protein structure by interrupting β-sheets.

**Figure 5.**
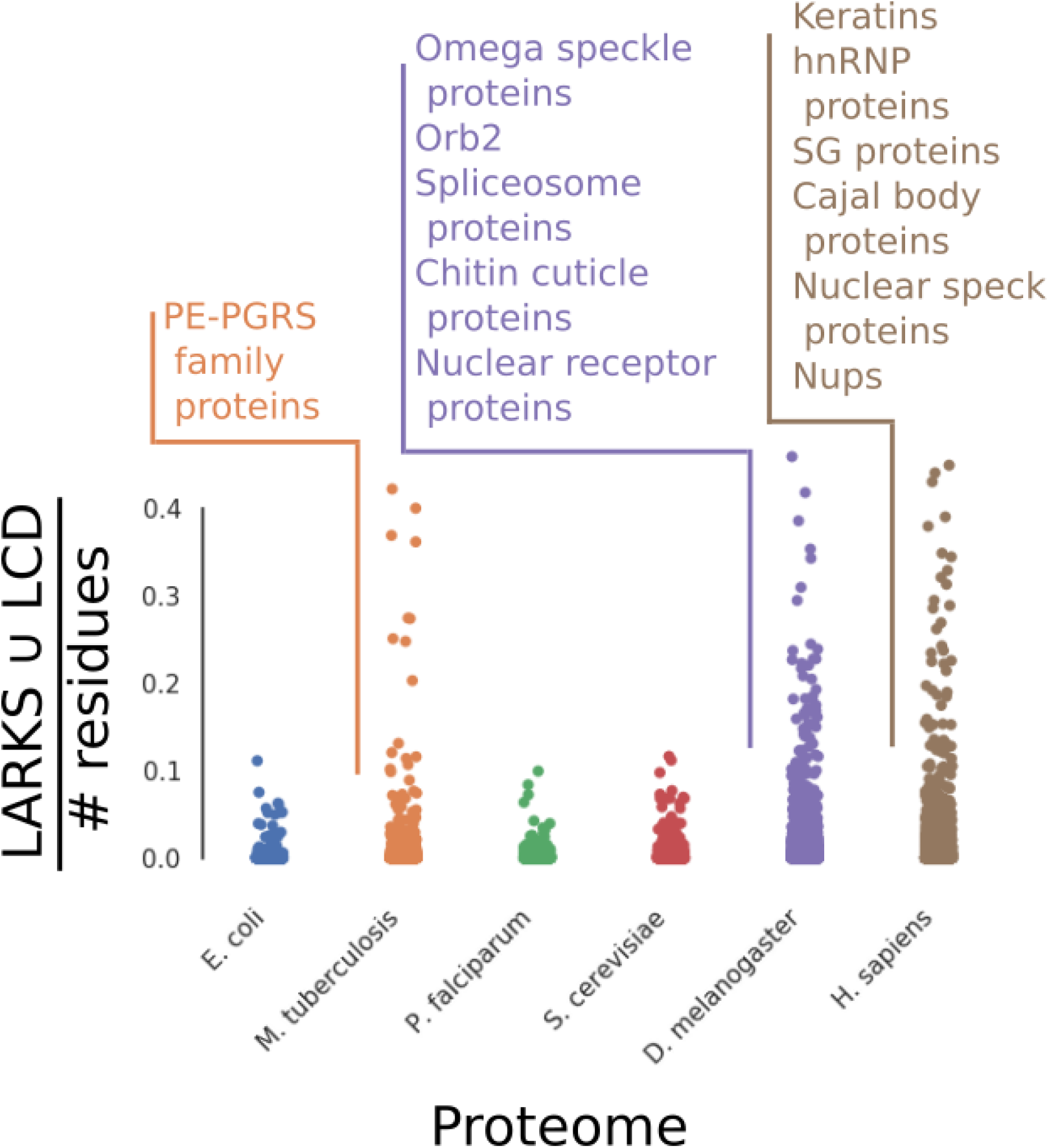
Swarm plot of LARKS∩LCD proteins in the six analyzed proteomes. Each dot represents a single protein and is placed on the Y-axis according to the number of LARKS∩LCD residues divided by the protein length. Examples of types of LARKS-rich proteins from *M. tuberculosis, D. melanogaster, and H. sapiens* are listed above and are largely associated with MLOs. Examples taken from Supplemental table 2.

*S. cerevisiae* does not show enrichment for LARKS∩LCD proteins in the proteome (Supplemental Figure 2), but when GO analysis is done on proteins with the most LARKS∩LCD residues, familiar GO terms are returned for mRNA binding, P-bodies, and stress granules (Figure 5; Supplemental Table 1). This finding appears to contrast slightly with our findings that LARKS-rich proteins do not significantly overlap with LCD-containing proteins in *S. cerevisiae* (Supplemental Figure 1). We believe discrepancy arises because while there are not many LARKS∩LCD residues in *S. cerevisiae*, looking for proteins with LARKS∩LCD residues finds proteins with LCDs, and GO terms may highlight an overlap of function of LCDs even if they are not LARKS-rich. If that is the case then the *S. cerevisiae* proteome does not enrich for LARKS in LCDs, but LCD containing proteins might serve similar functions for mediating phase-separation as in *H. sapiens* but using different motifs than LARKS. Yeast stress granules are less dynamic and more resemble aggregates than human stress granules (Kroschwald et al., 2015) which might stem from the LCDs in *S. cerevisiae* being more predisposed to forming steric zipper motifs over LARKS. So, while LCDs seem common on proteins involved in membraneless organelles, the LARKS content seem to tune material properties of condensates.

In *P. falciparum* a term for catalytic activity was found only in GO terms for LARKS-rich proteins, but we believe this may be spurious and reflects a few LCDs with a low number of LARKS within them, as *P. falciparum* has far fewer proteins with LARKS∩LCD residues (Figure 5). The proteome of *M. tuberculosis* has a population of proteins that are uniquely replete with LARKS∩LCD residues and returns unclassified GO terms. These proteins belong to the PE_PGRS family which is a mysterious family of proteins with Gly-Ala rich LCDs that decorate the outside of the cell wall (Brennan, 2017). In fact, Gly-Ala repeat inclusions are associated with C9orf72 related ALS and sequester cell machinery (Guo et al., 2018; Radwan et al., 2020). We find it striking that the LARKS∩LCD proteins on an obligate intracellular parasite are exposed to the host cell’s cytoplasm. It is enticing to speculate that these LARKS-rich proteins may have co-evolved with *H. sapiens* in order to interact with LARKS-rich proteins host cells to help virulence. *P. falciparum* is also an obligate intracellular parasite, but compared to *M. tuberculosis*, the proteome is largely devoid of LARKS in LCDs. In the *P. falciparum* life cycle, the parasite is endocytosed by a red blood cell. Once in the vacuole, the parasite modifies the environment within to better suit it but is not exposed to the intracellular environment. This contrasts with the *M. tuberculosis* parasite which can escape phagosomes to interact with the host cytoplasm (Russell, 2016). LARKS-rich proteins in humans are largely intracellular, involved in MLOs, and the LARKS-rich proteins displayed on the mycobacterium cell wall would be available to interact with the host machinery. Both parasites have extensive LCDs that must aide in virulence, but as their location and LARKS content is different their mechanism of actions is likely different as well.

Taken together, the variable LARKS content of LCDs may reflect the utility and physical behavior of that protein. For example, the LARKS-rich LCDs of human SG proteins seem to be more liquid-like than the LARKS-poor yeast stress granule proteins (Kroschwald et al., 2015), which behave more like solid aggregates. We reasoned that this difference in behavior could be reflected in structure, and to this end we made use of the recent surge of amyloid structures determined by cryoEM and ssNMR to interpret the effect that LARKS have on fibril structure. We opted to qualitatively compare the structures of putative functional amyloids of SG associated proteins FUS and hnRNPA2 that have numerous LARKS – 11 and 9 respectively – compared with the structure of Tau from disease-related amyloid that only has 2 LARKS (Figure 3). The Tau structure is a continuous β-sheet that folds onto itself to create an extended steric zipper that is the backbone of irreversible fibrils found in Alzheimer’s disease. The FUS and hnRNPA2 structures have a kinked backbone that interrupts the formation of any extended β-sheet, reflecting the number of LARKS in the structure.

How “kinked” a structure is can be roughly quantified by measuring the distance between Cα_i_ →Cα_i+4_. The pleated β-sheets in steric zippers have a consistent distance around 13.3Å, compared to LARKS where the kinks can reduce this distance to 7.5-12.6Å (Figure 4). The average Cα_i_ →Cα_i+4_ distances in FUS and hnRNPA2 structures are significantly shorter (10.8Å and 11.2Å respectively) than in Tau (12.4Å), consistent their respective LARKS content (Figure 3; Figure 4). The limited interfaces created by LARKS are associated with less stable amyloid-like fibrils (Hughes et al., 2018), fitting given the reversibility of FUS and hnRNPA2 fibrils (Kato et al., 2012; Murakami et al., 2015; Lu et al., 2020). This structural comparison of putatively functional, reversible amyloid fibrils hnRNPA2 and FUS (Kato & McKnight, 2018) to Tau support that LARKS alter structure in a way that is compatible with the biology of the different amyloid fibrils.

Not all functional amyloid is readily reversible, as in the case with Ordb2 from *D. melanogaster*. Orb2 forms a stable amyloid to aid in memory formation – stable amyloid is important to retain the memory over time (Hervas et al., 2020). The fibril-forming sequence of Orb2 contains a LCD that overrepresents glutamine residues, but has no predicted LARKS (Figure 3; Figure 4). The cryoEM structure of Orb2 amyloid fibril revealed a nearly perfect steric zipper of interdigitated glutamine residues. While a functional amyloid, Orb2 is better served by being irreversible and this behavior is reflected in the lack of LARKS in the Orb2 sequence.

This correlation of LARKS with kinked structures even holds within the same LCD. There are now two structures of the FUS LCD and four of the TDP43 LCD (Murray et al., 2017; Cao et al., 2019). The segments with more LARKS – PDBIDs: 5w3n and 6n3c - are more kinked than ones with fewer as judged by shorter Cα_i_ →Cα_i+4_ distances (Supplemental 3; Supplemental 4). FUS and other stress granule proteins form hydrogels that can be reversed several times before becoming irreversible (Murakami et al., 2015; Lu et al., 2020). It may be that the more kinked fibrils formed by LARKS may represent the labile form that may be a kinetic trap that ultimately yields to the more thermodynamically stable irreversible fibril core over time.

In summary, our findings support the hypothesis of LARKS as structural motifs in LCDs that can help mediate phase-transitions and MLO organization in biology. We find LARKS abundant in species that have dynamic MLOs (e.g. *D. melanogaster, H. sapiens*), depleted in species that have less dynamic stress granules (*S. cerevisiae*), and suspiciously abundant in intracellular parasites proteins that are exposed to the cytoplasm (*M. tuberculosis*). LARKS are not universal in all LCDs (e.g. *S. cerevisiae, P. falciparum*), and not in all proteins that undergo phase-transitions. LARKS are one strategy that organisms have adopted for organizing MLOs, but not all proteins that phase-separate need LARKS. Instead LARKS may tune the material properties of the condensates they are in by providing weak interactions for phase-separation while avoiding stable aggregation mediated by steric zippers. The degree to which beta-sheet structures form in MLOs remains debated (Peran & Mittag, 2020), but we show that amyloid-like structures rich in LARKS have kinked structures, and the kinked structures follow the principle of protein negative design by interrupting exposed β-edges, thereby preventing pathogenic aggregation in MLOs (Richardson & Richardson, 2002). LARKS appear to be a functional structural motif coopted through evolution by widely divergent species.

All the data presented in this work are available at LARKSdb. This permits researchers to submit their own proteins for LARKS predictions.

## Materials and methods

### Proteome Origin

All proteomes used were uniprot reference proteomes. The following proteomes and date of downloads were used: Escherichia coli: 2017-09-04, Mycobacterium tuberculosis: 216-04-04, Saccharomyces cerevisiae: 2016-04-04, Plasmodium falciparum: 2016-04-25, Drosophila melanogaster: 2016-04-04, and Homos sapiens: 2016-03-28. Any protein sequence with a letter not in the 20 natural amino acids was discarded before analysis.

### Identifying LARKS

LARKS were identified using the methods outlined in Hughes et al., 2018. The computational methods identify six-residue segments that are predicted to form LARKS segments. When counting the number of residues found in LARKS we considered any residue within the six-residue LARKS segment as a LARKS residue.

### Identifying LCDs

Residues in LCDs were identified by using the SEG algorithm (Wootton 1994). If 25 residues in a row were low in complexity as predicted by SEG, it was considered to be an LCD within a protein. Regions of low-complexity shorter than 25 residues were not considered to be in LCDs in our analysis.

### Identifying IDRs

IDRs were identified using Globplot 2.0 (Linding et al., 2003). The given script was downloaded and used to search proteomes for predicted globular regions. Any residue predicted to be in a globular region using the standard settings recommended was considered to globular, and all other residues predicted to be in IDRs in our analysis. Computational overlap of predicted residues

All residues were given binary scores (either belonging to a category, or not) in each category: LARKS, LCDs, IDRs, and steric zipper (steric zippers analysis only for a small number of proteins in Supplemental Figure 1). Then residues in overlapping categories could be identified. Enrichment of overlapping classes of residues (e.g. LARKS in LCDs) was calculated by finding the actual number of residues predicted to be both LARKS and in LCDs within a proteome and comparing this to an expected number. The expected overlap was found for each proteome by multiplying the probability of residues in one category (e.g. LARKS) by the probability of a residues being in another category (e.g. LCDs).

### Bootstrapping methodology

For each proteome computed the average number of LARKS per 100 residues across the proteome. Any protein that has more than the average (given in the table below) for the organism was considered LARKS-rich.

| Species | Average # LARKS per 100 residues |
| --- | --- |
| *E. coli* | 2.2 |
| *M. tuberculosis* | 3.1 |
| *P. falciparum* | 0.9 |
| *S. cerevisiae* | 1.7 |
| *D. melanogaster* | 2.2 |
| *H. sapiens* | 2.2 |

The number of proteins with an LCD and considered LARKS-rich was counted for each proteome to give the actual value (Figure 1). Then bootstrapping was done by randomly drawing proteins, with replacement, from the proteome equal to the actual number of proteins with LCDs that were counted from that proteome. From that sample, the number of LARKS∩LCD proteins were counted. This process was repeated 10,000 times to build a distribution of numbers of LARKS∩LCD proteins that are found in the random samples of the proteome equal in size to the number of LCD proteins. The actual number of LARKS∩LCD proteins was counted and plotted on the histogram to see if it was above, below, or within the distribution – actual values given in the table below (Supplemental figure 2). A p-value was calculated by counting the number of random samples that had values that exceeded the actual number of LARKS∩LCD proteins and dividing by the number of samples (except for P. falciparum where the number of samples with fewer LARKS∩LCD proteins to find depletion of LARKS∩LCD in LCD proteins).

| Species | # of samples > LARKS∩LCD actual | # of samples < LARKS∩LCD actual |
| --- | --- | --- |
| *E. coli* | 3 | 9997 |
| *M. tuberculosis* | 0 | 10000 |
| *P. falciparum* | 10000 | 0 |
| *S. cerevisiae* | 2884 | 7116 |
| *D. melanogaster* | 0 | 10000 |
| *H. sapiens* | 0 | 10000 |

### Gene Ontology

GO was done on the top 5% of LARKS-rich proteins in each proteome with LARKS-rich defined as the average number of LARKS per 100 residues of a protein. The list of uniport IDs for these top 5% of proteins was submitted to the Panther GO server to see enriched terms (Thomas et al., 2003). Relevant GO terms were selected arbitrarily for Figure 5 and the complete list of GO term hits is in supplemental table 1.

## Abbreviations

LARKS: Low Complexity, Amyloid-like Reversible Kinked Segments
LCDs: Low-Complexity Domains
PrLDs: Prion-like Domains
IDRs: Intrinsically Disordered Regions
LLPS: Liquid-Liquid Phase Separation
MLOs: Membraneless Organelles
LARKS∩LCD protein: protein that has at least one LCD and for which the fraction of LARKS is greater than average for its proteome
LARKS∩LCD residue: an amino acid residue that is a predicted LARKS within a LCD

---

**Supplemental Figure 1.**
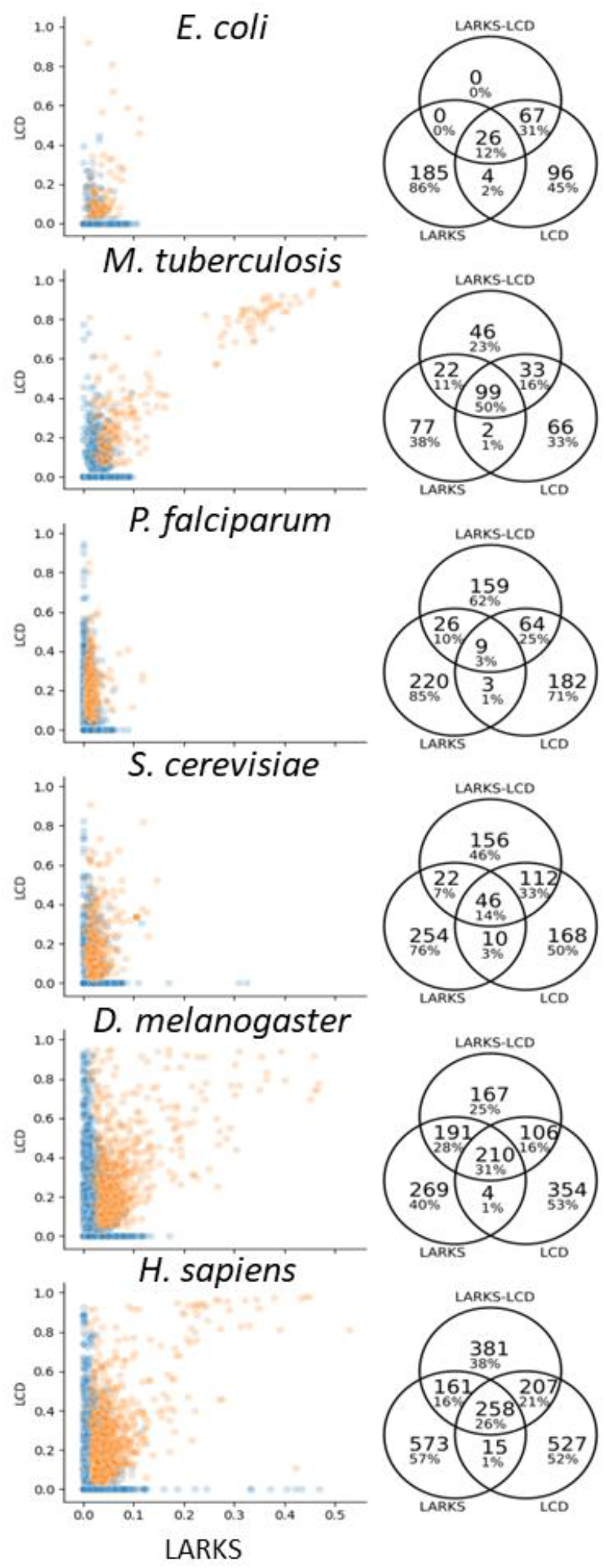
Proteins in proteomes plotted by fraction of residues in LARKS for the X-axis and fraction of residues in LCDs for the Y-axis. Each point represents a single protein. Proteins that are in the 95^th^ percentile of LARKS-LCD rich residues in each proteome are shown in orange, and GO analysis was performed on these top 5% of proteins to see enrichment in different functions of these proteins (Figure 5; Supplemental table 1). To show overlap of proteins in the 95^th^ percentile for most abundant residues in LARKS, LCDs, or LARKS-LCD we show overlap of these three groups in the Venn diagrams.

**Supplemental Figure 2.**
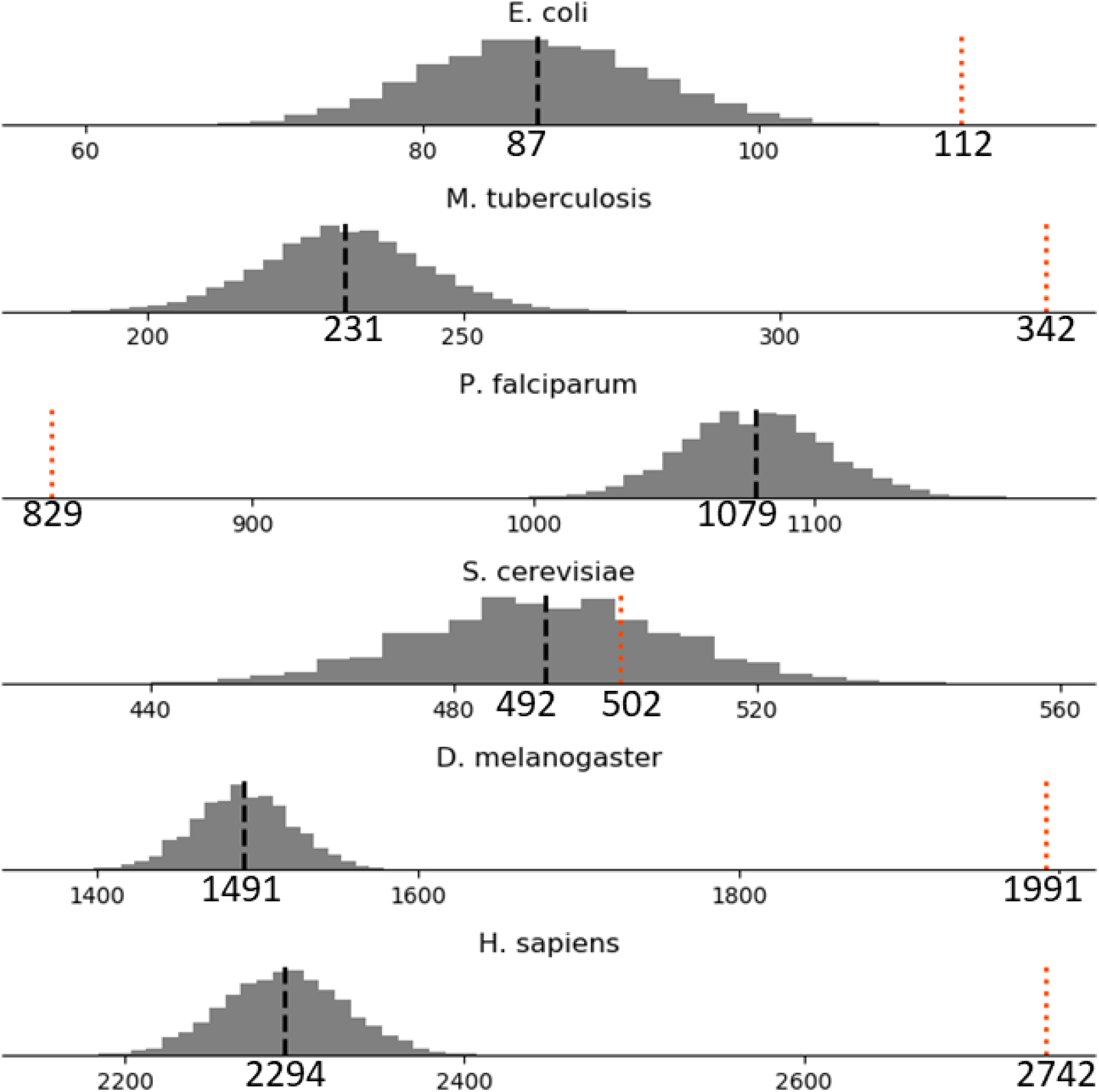
Bootstrapping statistics show LARKS-rich and LCD-containing proteins significantly overlap in some organisms. For each proteome shown in Figure 1, the number of LARKS∩LCD proteins are shown by the red line. Then we drew proteins randomly with replacement from the species proteome equal to the number of proteins with LCDs in the proteome. From that random sample the number of LARKS∩LCD proteins that had an LCD and were LARKS-rich were counted, and this process was repeated 10,000 times to create a distribution of averages. The distribution average is given by the black line and the actual average of LARKS∩LCD below it. The data indicate that LARKS are significantly enriched in LCD proteins for all species except *and S. cerevisiae*. In *S. cerevisiae* the actual number of LARKS∩LCD proteins is well within the expected distribution generated from bootstrapping, and in *P. falciparum* proteins considered LARKS-rich are less likely to have and LCD than expected by random chance.

**Supplemental Figure 3.**
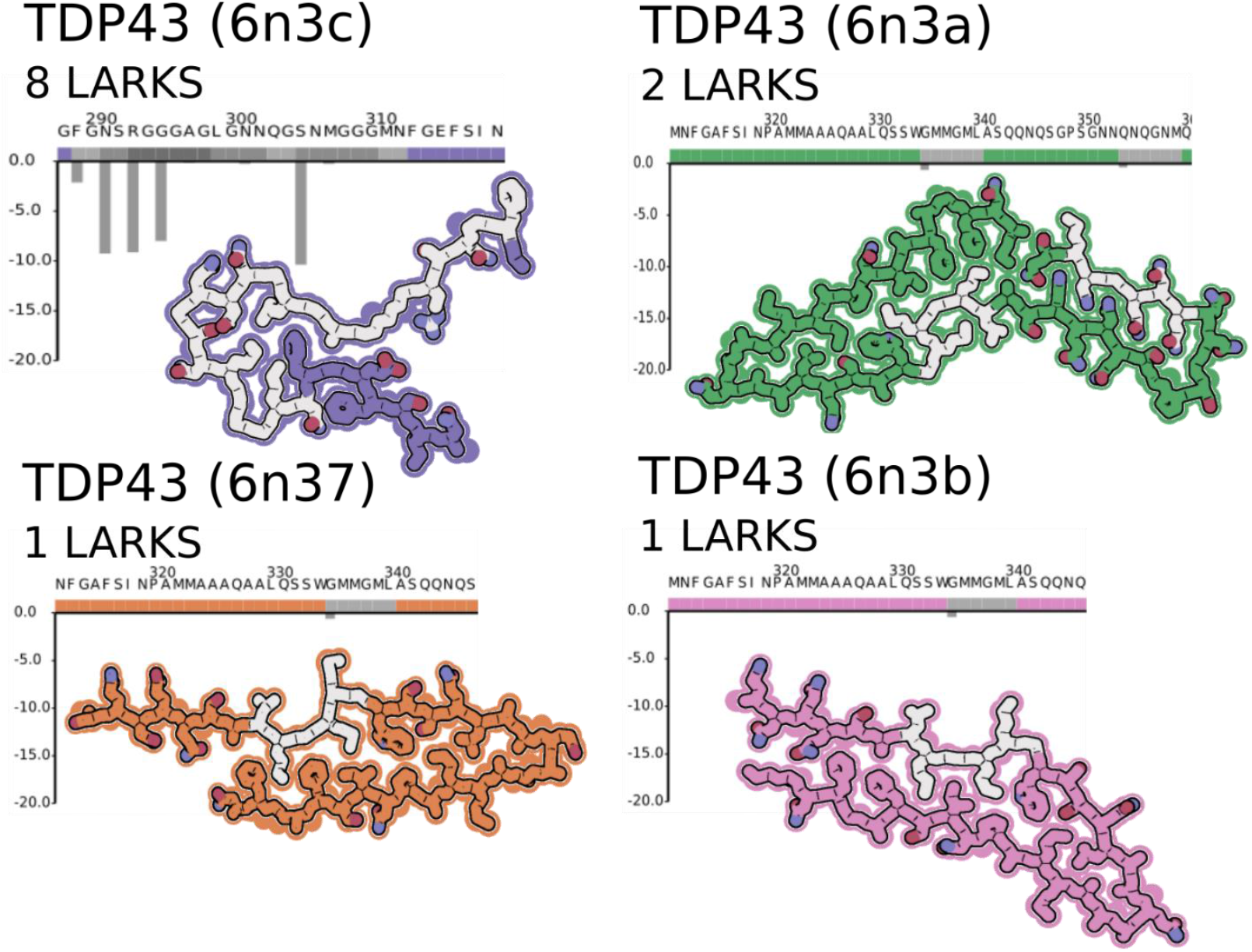
Three additional structures of polymorphic amyloid fibrils TDP43 from different regions of the LCD (PDBIDs: 6n37, 6n3a, and 6n3b) with few LARKS compared to a LARKS-rich region of the LCD (6n3c) to highlight the kinked backbone created by LARKS in 6n3c.

**Supplemental Figure 4.**
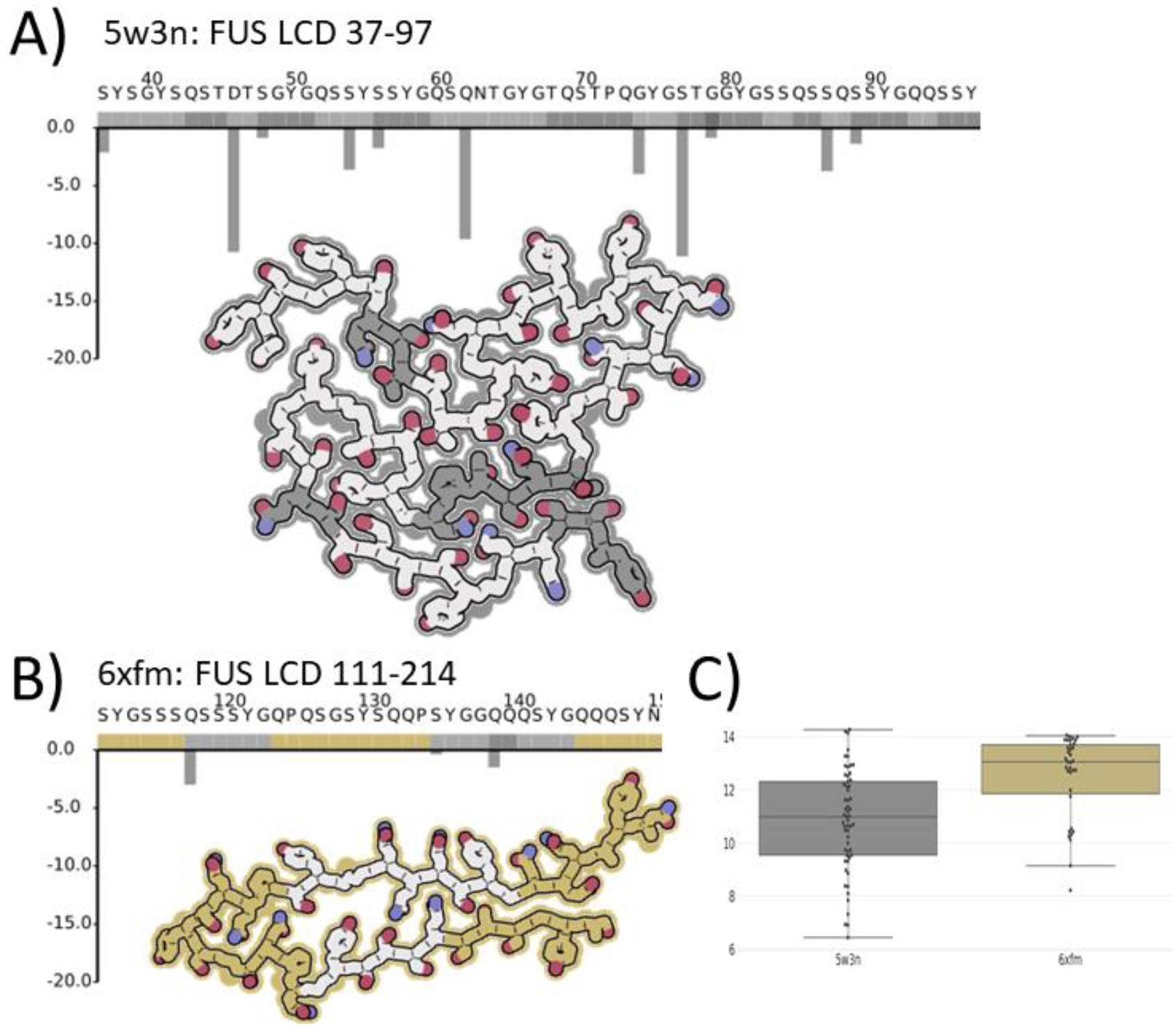
FUS LCD amyloid fibril structure comparisons. A) The FUS structure 5w3n, determined by ssNMR, consists of LCD residues 37-97. This segment contains 11 LARKS and a dramatically kinked backbone. B) The FUS structure 6xfm, determined by cryoEM, consists of residues 111-150, but only has 3 LARKS. C) Box plot of Cα_i_ →Cα_i+4_ distances from FUS structures shows that the average distance is shorter for the LARKS-rich 5w3n structure compared to the LARKS-poor 6xfm structure reflecting the effect of LARKS on the amyloid backbone.

